# Crystal Toxins and the volunteer’s dilemma in bacteria

**DOI:** 10.1101/437913

**Keywords:** virulence, cooperation, game theory, kin-selection, evolution, social evolution

## Abstract

The growth and virulence of the bacteria *Bacillus thuringiensis* depends on the production of Cry toxins, which are used to perforate the gut of its host. Successful invasion of the host relies on producing a threshold amount of toxin, after which there is no benefit from producing more toxin. Consequently, the production of Cry toxin appears to be a different type of social problem compared with the public goods scenarios that bacteria often encounter. We show that selection for toxin production is a volunteer’s dilemma. We make the specific predictions that: (1) selection for toxin production depends upon an interplay between the number of bacterial cells that each host ingests, and the genetic relatedness between those cells; (2) cheats that do not produce toxin gain an advantage when at low frequencies, and at high bacterial density, allowing them to be maintained in a population alongside toxin producing cells. More generally, our results emphasise the diversity of the social games that bacteria play.

## 2 Introduction

The growth and virulence of many bacteria depends upon successfully cooperating in public goods games with other bacteria. Bacteria produce and secrete a range of molecules, which provide a benefit to the local group of cells, and so act as public goods. For example, iron scavenging siderophores, or protein digesting proteases (West et al., 2007). Individual cells pay the metabolic cost of producing these molecules, but their benefits are then shared as public goods with the local population of cells. Consequently, producing cells could potentially be out-competed by non-producing, cheats, who gain the benefits, without paying the costs. There is a large theoretical and empirical literature examining how various factors such as interactions between genetically identical cells (kin selection), can stabilise the production of public goods in bacteria (Brown and Johnstone, 2001; West and Buckling, 2003; Griffin et al., 2004; Diggle et al., 2007; Frank, 2010*b*,*a*).

In contrast, the growth and virulence of the bacteria *Bacillus thuringiensis* appears to depend upon a different type of social game (Raymond et al., 2012). The life cycle of this bacteria depends upon two steps in the host. First, after an insect host ingests a number of spores, the bacterial cells use a costly crystal (Cry) toxin to perforate the host gut, and invade the host (Höfte and Whiteley, 1989; Ibrahim et al., 2010; Raymond et al., 2012). The toxin is a large protein, up to 147 kilodaltons, that may form up to 35% of a bacteria’s dry mass (Loferer-Krößbacher et al., 1998). Second, the bacteria multiply within the host and invest in Cry toxin production, causing host death and the release of bacterial spores (Raymond et al., 2010). In contrast to a public goods scenario the benefit of producing Cry toxin is all or nothing — you either produce enough to invade the host, or you do not. As producing a certain total amount of toxin is key, the strategy that will be favoured by evolution could also depend upon the number of spores that are inside a host (Archetti, 2009; Raymond and Bonsall, 2013; Cornforth et al., 2015).

We examine the evolutionary stability and dynamics of Cry toxin production using two different modelling approaches. First, we use a game theoretic approach to examine under what conditions the production of Cry toxin is favoured (Taylor and Frank, 1996). This approach assumes only small variations in toxin production (weak selection), and looks for a single equilibrium. In contrast, in nature there is large variation in toxin production, between cells that produce (cooperators) or do not produce (cheats) Cry toxin (Raymond et al., 2010, 2012; Deng et al., 2015). Furthermore, factors such as population density and cooperator frequency can fluctuate over short timescales (Schoener, 2011; Raymond et al., 2012; Gokhale and Hauert, 2016), and studies of the density of spores in the wild have shown that group sizes are very low suggesting that stochastic effects could be important (Maduell et al., 2002; Collier et al., 2005; Raymond et al., 2010). Therefore, our second approach is to model the dynamics of a system that contains both co-operators and cheats, to examine how these dynamics are influenced by bacterial density, and the frequency of cooperators.

## 3 Model I: Equilibrium Model

We use a game theoretic approach to express the fitness of a bacterial cell as a function of: the probability it infects a host, *β*(*z*); and, the number of spores it generates, *f* (*y*) — where, *z* is the group average strategy and *y* is the individual cells strategy. We assume an infinitely sized population of bacteria distributed into finitely sized patches of *n* bacteria. There are non-overlapping generations and the bacterial spores disperse randomly to other patches.

We assume that the probability that a bacteria in a group of *n* cells successfully infects a host, *β*, is a function of their average investment, *z*. We model this probability using a sigmoidal curve as a continuously differentiable approximation of a step function:

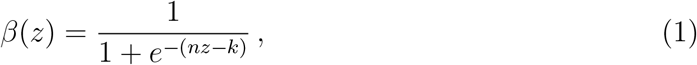

where, the group production of toxin *nz* is compared to *k*, which is the threshold at which the chance of infection would be 0.5 (Cornforth et al., 2012). When the total toxin production is low (*nz* << *k*) then the chance of infection is close to 0 as toxin production increases 0 ≤ *nz* ≤ *k* then the function is accelerating and then past the threshold (*k* < *cz*) the function is decelerating and asymptotes to 1.

We assume there is a linear trade-off between the energy a bacteria puts into producing toxin, *y*, and the energy available for growth, *f* — as both require the generation of protein:

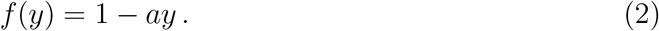

The fitness function of a focal bacterium will be the product of the probability it invades a host and the growth of the bacterium once it has successfully invaded (*β*(*z*) *· f* (*y*)):

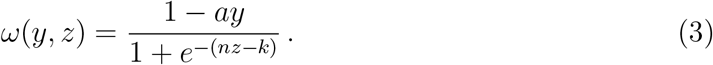

Equation (3) illustrates that producing the Cry toxin has a cost to the individual by reducing its growth, *f* (*y*). However, it is beneficial to the group, including our focal individual, as it increases the chance of successful invasion, *β*(*z*).

We seek an evolutionarily stable strategy (ESS), which is the individual strategy at fixation which cannot be invaded by some rare alternative strategy. Following Taylor and Frank (1996), we construct an expression for the change in inclusive fitness, ∆*ω_IF_*, and solve for a monomorphic population that is at equilibrium:

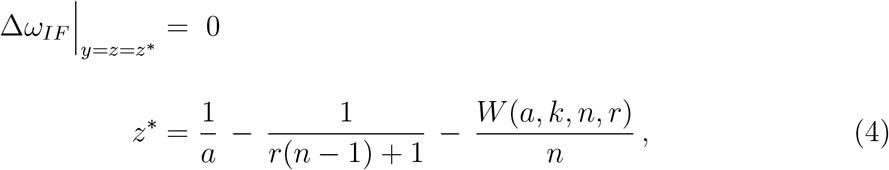

where, *W* is a Lambert-W function which is strictly positive (see B) and *r* is the relatedness between the different bacterial cells infecting the host. We define *r* as the probability that two individuals share the same gene at a locus relative to the population average (Grafen, 1985). This measure is obtained by replacing the regression of the recipients phenotype on the focal individuals genotype (*R* in Taylor and Frank (1996)) with: 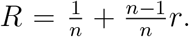 Where 1*/n* is the chance the other individual is oneself and *n* − 1*/n* is the chance of a social partner with other’s only relatedness *r* to the focal individual (Pepper, 2000).

The equilibrium at *z*^*^ is a maximum however it may be unreachable. To test whether a population under weak selection would converge to equilibrium (convergence stability), we examined whether the second order terms at the equilibrium were negative (Otto and Day, 2011). We found that:

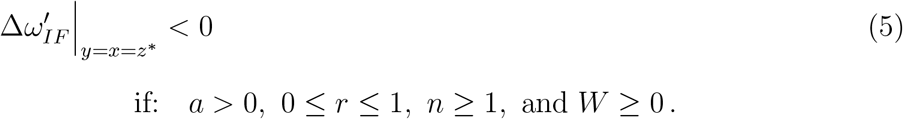

So the equilibrium at *z*^*^ is a candidate ESS. To determine uninvadibility we implement an extension to the Taylor and Frank (1996) approach, by interpreting the second derivative of the fitness equation in terms of inclusive fitness effects, therefore establishing a condition for the candidate equilibrium to be a local maximum (Cooper and West, 2018). In A we show that *z*^*^ is an uninvaidable strategy as well as being convergently stable.

### 3.1 The effect of relatedness

We found that increasing relatedness (*r*) increases individual toxin production. Examining the derivative of the equilibrium toxin production (*z*^*^) with respect to relatedness (*r*) we found that:

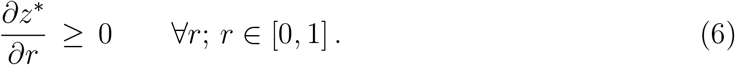

So as relatedness (*r*) within the group increases the ESS of toxin also increases (*z*^*^) (C). Increasing relatedness increases the indirect benefit from toxin production as the group chance of invasion, *β*(*z*), has a greater chance of being shared with kin. However, even when relatedness is low (*r* = 0) toxin production is favoured as it is essential to reproductive success (fig. 1).

**Figure 1:**
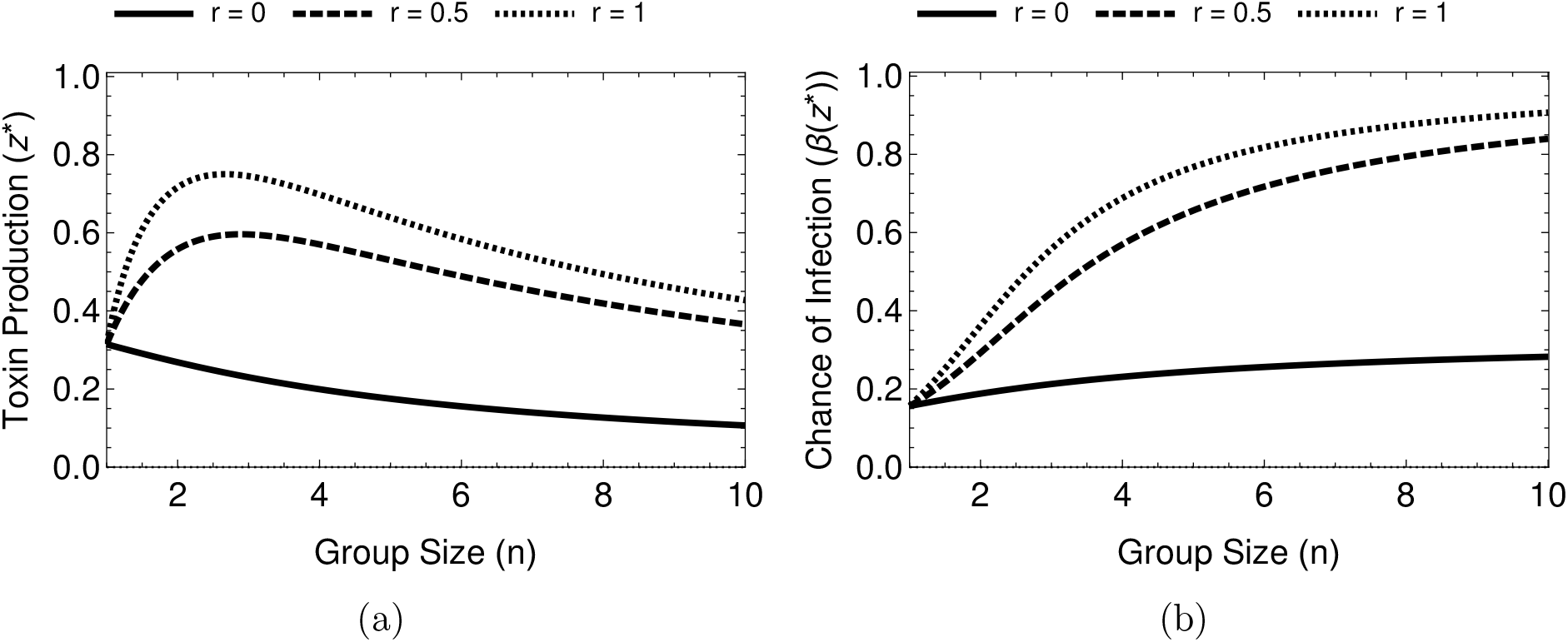
The equilibrium toxin production depends on group size (*n*) and relatedness (*r*). **a)** When *r* > 0, as we increase group size, toxin production initially increases and then decreases. **b)** The total amount of toxin produced by the group, *nz\**, increases with group size, therefore, the chance of infecting the host is always higher in larger groups. These graphs assume *k* = 2 and 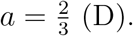

### 3.2 The effect of group size

As groups increase in size individual toxin production initially peaks and then declines — when relatedness is non-zero (fig. 1a). This is due to the efficiency gained when close to the accelerating section of the sigmoidal *β*(*z*) function (near the threshold). As the benefits (*β*(*z*)) are accelerating, small increases in toxin production lead to large increases in infection chance. Past the peak toxin production, the greater number of individuals in the patch allow for individual bacteria to reduce their investment but the group remains at a high chance of successfully invading (see D).

### 3.3 The effect of the threshold

The derivative of toxin production, *z*^*^, with respect to the threshold is always positive or zero:

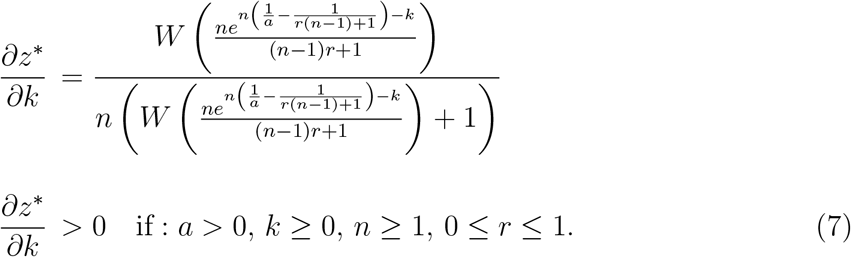

Therefore, in the absence of other limits if more toxin is required to invade the host (higher *k*) individuals will be selected to increase their toxin production (*z*^*^).

### 3.4 Cry toxin as a Volunteer’s Dilemma

Our model illustrates that the production of crystal toxin by the bacteria is a volunteer’s dilemma. Volunteer’s dilemmas are a class of social games where the benefit is gained after a threshold investment in the good is reached and the benefit is fixed for each member regardless of group size or personal investment (Archetti, 2009, 2018). The perforation caused by Cry toxin can be used by any organisms in the gut, it is a public good. The Cry toxin only perforates the gut after a certain concentration, the good acts after a threshold (eq. (1)). And, the benefit to the bacteria is access to the tissue of the host which is a binary outcome, either the bacteria have access or not, there is no additional benefit for exceeding the threshold of Cry toxin in the gut (Höfte and Whiteley, 1989; Ibrahim et al., 2010; Raymond et al., 2012). These qualities of the Cry toxin system make its production a volunteer’s dilemma.

## 4 Model II: Cooperator-Cheat Dynamics

In nature the density and fraction of spores, that either do (cooperators) or don’t (cheats) produce Cry toxins, can be very variable over short temporal and spatial scales (Maduell et al., 2002; Collier et al., 2005; Raymond and Bonsall, 2013). We capture this ecological variation with a model which allows us to compare individuals that produce toxin at a fixed level (cooperator) against individuals which do not produce any toxin (cheats). We compare the relative fitness between these two types to determine under varying ecological parameters.

We assume a population of bacteria whose spores freely mix and are taken up at random by a host. We assume that the host ingests *P* bacterial spores. In the environment a proportion (*c*) of bacteria are cooperators and (1 − *c*) are cheats. For a focal individual in a group of *P −* 1 social partners there are *i* cooperators which are distributed:

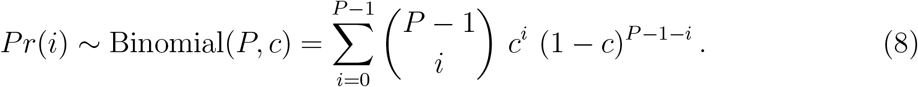

From eq. (3) given *i* cooperators in a group the payoff, *π*, for the focal bacteria producing *y* toxin will be:

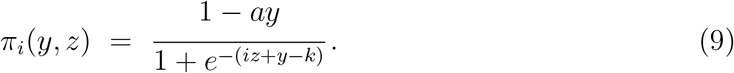

Therefore, the overall fitness of a focal bacteria producing *y* toxin in a population of cooperators producing *z* toxin will be:

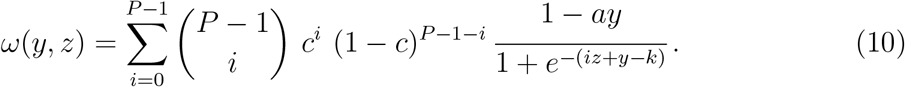

This allows us to express the fitness of a cooperator in the population as *ω*(*z, z*) and that of a cheat as *ω*(0, *z*). The relative fitness of cheats to cooperators in the population is:

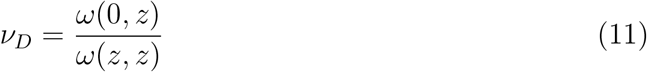

### 4.1 Frequency dependence

As the proportion of cheats increases we find that the relative fitness of cheats decreases (fig. 2a). As cheats become more common, groups become dominated by cheats and the chance that a group produces enough toxin decreases. Why do we find frequency dependence when, in the simplest possible case, a public goods dilemma leads to selection being frequency independent (Ross-Gillespie et al., 2007)?

**Figure 2:**
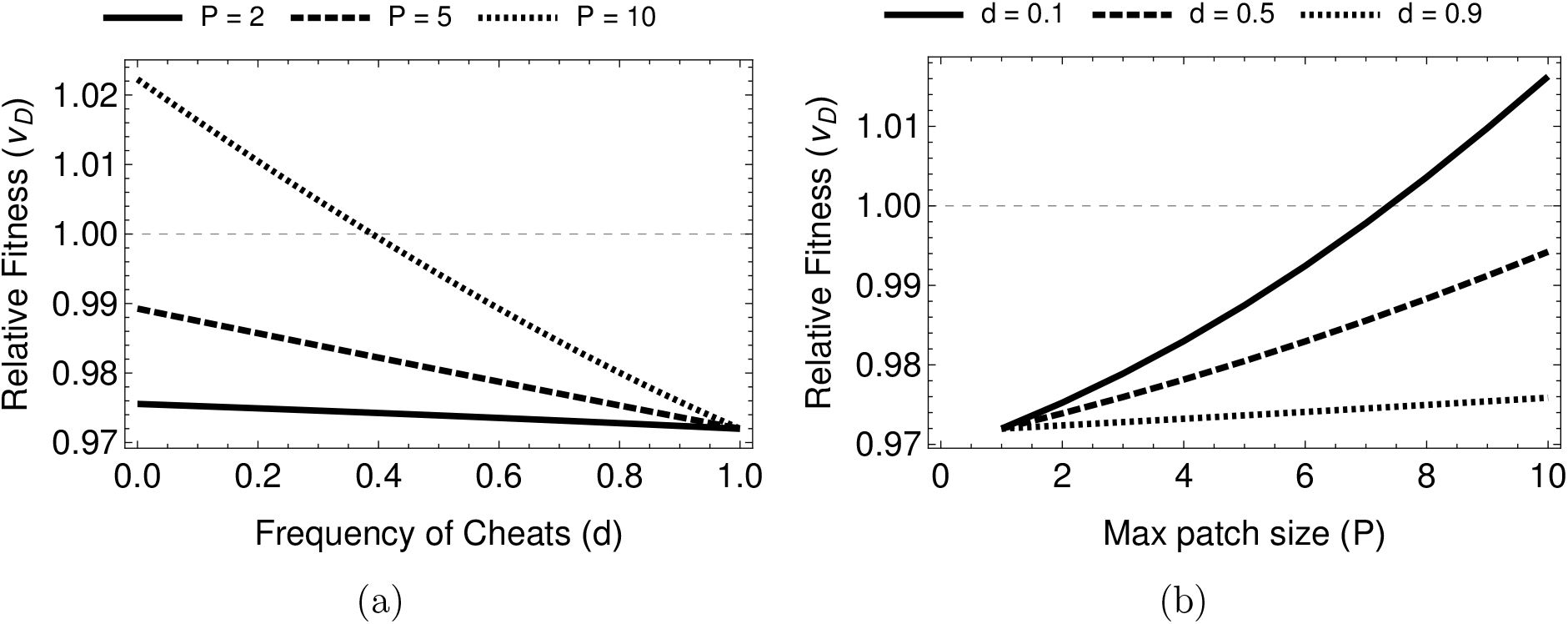
**(a)** The relative fitness of cheats is negatively frequency dependent as cheats become more common they are more often aggregated together and so suffer in relative fitness to cooperators. **(b)** As group size increases there is a positive density dependent effect on cheat fitness, the larger the group the more chance that sufficient toxin is produced by the group

The result of frequency independence requires either: (1) that the effect on public good production is linear or, (2) that the trait is under weak selection. Either of these two assumptions make a linear approximation, using a first order Taylor expansion, valid. And, such expansions, are frequency independent (Rousset, 2004; Lehmann and Rousset, 2014). This argument is similar to the justification for frequency independent selection of a trait that the selection gradient, *s*(*z*) = *∂ω/∂y* + *r∂ω/∂z*, is constant with respect to allele frequency (Hamilton, 1964; Gore et al., 2009; Lehmann and Rousset, 2014).

However, in our model we find that the relative fitness of a cheat is frequency dependent. This is because we relax both of the assumptions made by Ross-Gillespie et al. (2007). We have a non-linear synergistic effect between cooperators which means that each cooperator or defector does not have a linear effect on the fitness of the focal individual, due to the step like benefit function (*β*(*z*)). Addition or subtraction of a cooperator has a large effect when a group is close to the the threshold but a much smaller effect when the group toxin production is already very low or very high; the benefit of a cooperator is dependent on the compisition of the group which is itself dependent on the frequency of cooperators. This synergy introduces a frequency dependent term into the first order effects of our selection gradient (Lehmann and Rousset, 2014). Secondly, we consider a game with strong selection which makes approximating the gradient using only first order terms inappropriate. The large difference between cooperator and cheat strategies causes higher order terms of the relative fitness to matter and these higher order terms will include frequency dependent terms (Hamilton, 1964; Ross-Gillespie et al., 2007).

These two effects lead to a frequency dependent relative fitness found here — unlike the frequency independence found in earlier models (Ross-Gillespie et al., 2007). The synergistic game causes the first order term of the Taylor expansion to be frequency dependent. The strong selection causes higher order terms to become more substantial. These two effects are sufficient but not necessary conditions for frequency dependence to arise.

### 4.2 Density dependence

Increasing the density of the population, by increasing the group size (*P*), increases relative cheat fitness. In more dense populations there is a greater chance that a group will have a sufficient number of cooperators to invade a host successfully (fig. 2b). The mean number of cooperators in a group increases with density allowing cheats to exploit more cooperators. In the limit, as *P* increases, the chance of infection for all patches in the population is one, (*β*(*z^*^*) = 1). Therefore, the fitness of cheats is 1 and the fitness of all cooperators is 1 − *az*. The relative fitness of cheats then approaches, 1/(1 − *az*).

### 4.3 Population Aggregation

The above model assumes patches form randomly from the population with no structuring beyond random chance. We now imagine a scenario where similar strategies are clumped together, as would be expected if they had emerged from the same host (van Leeuwen et al., 2015). We use a modified Poisson binomial distribution to model the initial member of a group biasing subsequent draws towards its own type. As initial founders are randomly distributed, a fraction *c* of the groups are clumped around a cooperator and a fraction *d* around a cheat (*c* + *d* = 1).

So given that the patch is started by a cooperator then the distribution of number of cooperators among such patches is:

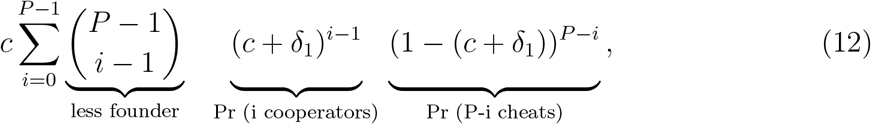

and similarly for cheats:

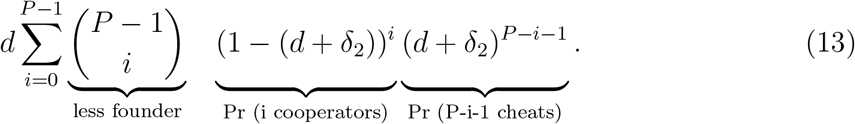

The binomial coefficient is *C*(*P −* 1, *i* − 1) for cooperators as the founder individual counts for the first group member and the first cooperator. For the defector patches the founder only accounts for the first group member, hence *C*(*P −* 1, *i*). The two variables, *δ*_1_ and *δ*_2_, are terms that bias the distributions based on the founder. The larger their values the more strongly the two types aggregate. These two distributions represent an underlying distribution — that of the simpler model. We define *ϕ* ∈ [0, 1] as the level of aggregation and define the bias parameters as: 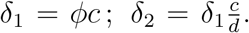 When *ϕ* is one then patches of all cooperators and all cheats form and when it is zero then there is no bias and patches form as they would in a binomial distribution. By expressing the bias parameters (*δ*_1_ and *δ*_2_), in terms of *ϕ*, *c* and *d*, we ensure that the sum over both distributions is equal to one, and the terms are weighted probabilities.

The distribution of the number of cooperators in a patch is weighted by the fitness of the focal individual in such a group (the sum of the above two distributions), giving:

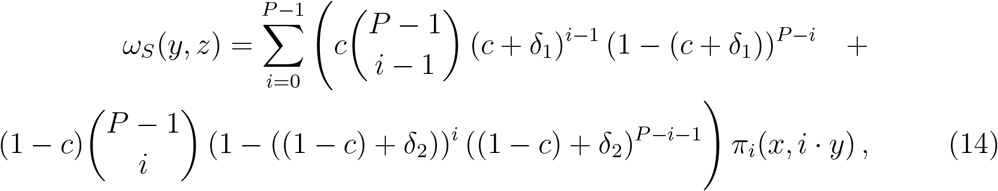

and from this we calculate a structured relative fitness: *ν_DS_* = *ω_S_*(0, *z*)*/ω_S_*(*z, z*)

At maximum aggregation (*ϕ* = 1) cheats will do very poorly against cooperators as groups formed of all cheats have almost zero chance of invading the host. In the absence of aggregation (*ϕ* = 0), cheats will be performing as if the population were unstructured, as in the previous model. As aggregation increases cheats are more likely to find themselves in groups composed mostly of cooperators or mostly of cheats and very rarely a group close to an unbiased distribution.

Intermediate levels of aggregation can, with intermediate frequencies of cooperators and high densities, lead to an increase in cheat relative fitness (fig. 3). When group sizes are large the benefit to all members of a group of cooperators will approach one. At that point any additional cooperators will perform much worse than additional cheats as they will be paying the cost of producing toxin and gaining no marginal benefit from this additional toxin (infection chance can’t be greater than one). Therefore, at intermediate levels of aggregation, enough cooperators will be on patches to infect a host and be exploited by cheats. Conversely in the defector biased groups the threshold will never be reached and cooperators perform poorly as they are paying a cost for little benefit and any generated benefit is being exploited by cheats. This leads to high density scenarios with intermediate levels of aggregation increasing cheat relative fitness.

**Figure 3:**
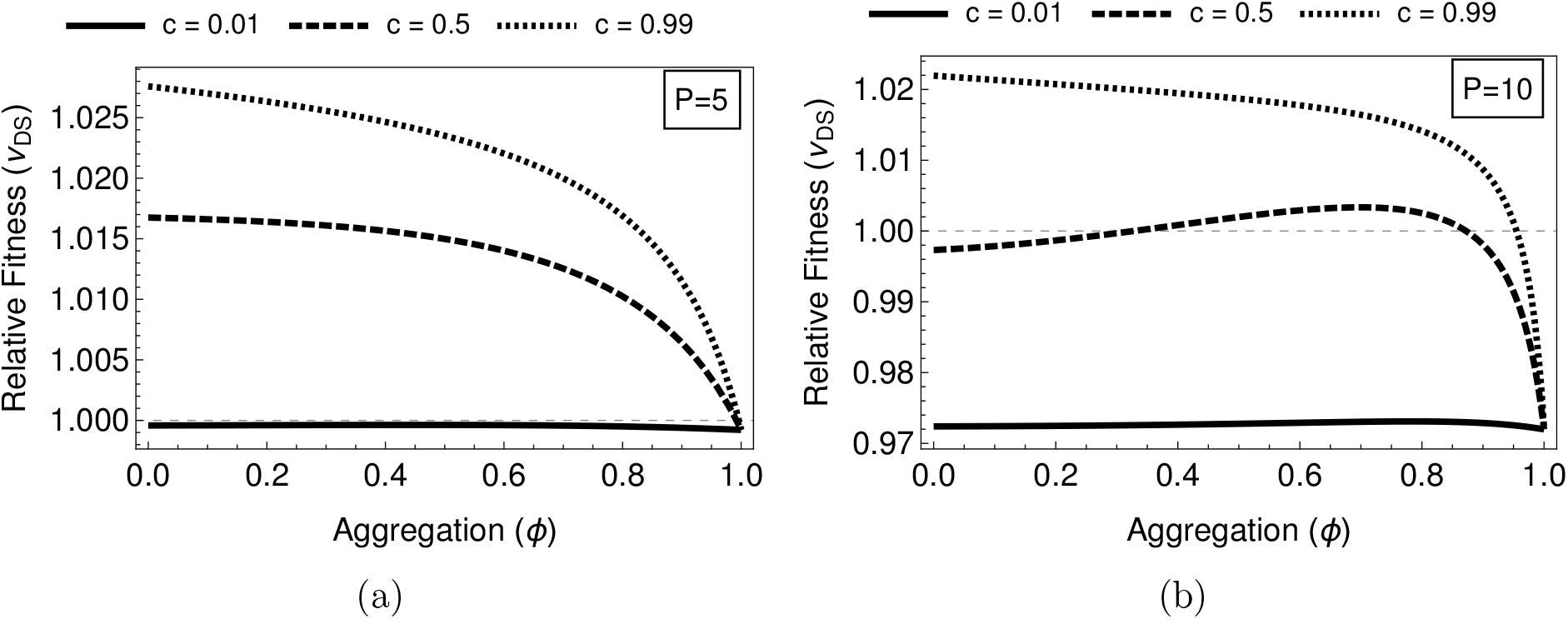
Graphs of *ω_S_*, eq. (14), using parameters: *k* = 2, *a* = 2/3 and *z* = 0.17. **(a)** When group size is low, *P* = 5 increasing aggregation leads to decreasing relative fitness for cheats regardless of the initial cooperator frequency **(b)** At higher group sizes (*P* = 10) the pattern is also decreasing at high cooperator frequencies however at middling and low densities we see a non monotonic pattern with an intermediate aggregation causing a maximum relative fitness in cheats.

The above method of looking at the relative fitness of cheats to cooperator shows whether a cheat will be increasing or decreasing in the population. This gives a static view of the dynamics occurring in the population. Our analysis shows how cheats can have a high enough relative fitness to invade a population and some predictions on what would happen as environmental and demographic parameters change. However, they cannot establish over the long term whether cheating is a stable strategy in a population. So, in E we show that cheat-cooperator co-existence can be reached dynamically from our model (Peña et al., 2014; Archetti, 2018).

## 5 Discussion

We found that the production of toxin by the bacteria *B. thurigiensis* is diferent from classical public goods games. The threshold nature of the toxin production leads to a volunteer’s dilemma where for each individual it would be optimal if another were to volunteer to produce the good instead of them. We found, with a game theory approach, that the ESS level of toxin production: (1) increases when the cells infecting a host host are more related, and (2) peaks at intermediate numbers of cells infecting a host (fig. 1). We then developed a stochastic model of the dynamics of cooperators that produce toxin, and cheats that do not produce toxin. We found the relative fitness of cheats was greater when: (1) they were less common (lower frequencies), (2) more cells infect each host (higher densities) (fig. 2), (3) cells tended to be aggregated with the same cell types (relatives) (fig. 3). Our results show how ecological conditions can influence the relative fitness of cheats and cooperators, in ways that could feedback into the population dynamics of *B. thurigiensis* and its invertebrate hosts.

### 5.1 The volunteer’s dilemma for public goods

Our finding that toxin production resembles a volunteer’s dilemma game leads to some different predictions compared with most other social traits in bacteria, which represent public goods games (Brown, 1999; Brown and Johnstone, 2001; West and Buckling, 2003; Ross-Gillespie et al., 2007; Ross-Gillespie et al., 2009; Frank, 2010*a*). We found that individual investment (toxin production) is highest at intermediate group sizes, that the fitness of cheats can depend upon their frequency in the population (frequency dependence) in well mixed populations, and that intermediate levels of aggregation can increase the relative fitness of cheats (Archetti, 2009; dos Santos and Peña, 2017; Archetti, 2018). In contrast, in public goods games, toxin production is not frequency dependent in well mixed populations, and intermediate levels of aggregation decrease the relative fitness of cheats (Brown and Johnstone, 2001; West and Buckling, 2003; Ross-Gillespie et al., 2007; Ross-Gillespie et al., 2009).

Our result that cheater fitness is dependent upon the frequency in the population contrasts with Hamilton (1964)”gift from god” that cooperator fitness should be independent of frequency. Our analyses differ from Hamilton’s in two ways. Firstly, in the volunteer’s dilemma, each additional player has a non-linear effect (non-additivity) on the benefit, which means that even when looking at first order terms frequency is present as a variable (Rousset, 2004). Secondly, in our models we assume that the cheater produces no toxin and the cooperator produces a large quantity, leading to strong selection, which means that linearising the relative fitness is no longer appropriate as higher order terms have large effects (Ross-Gillespie et al., 2007; Gore et al., 2009; Lehmann and Rousset, 2014).

### 5.2 Bt in the wild

Our results are supported by both observational and experimental data from field populations of *B. thurigiensis*. Consistent with our prediction that frequency dependent selection can lead to to cooperators and cheats coexisting, natural populations show variation in the level of cry toxin production, with both producers and non-producers coexisting (Raymond et al., 2010, 2012; Raymond and Bonsall, 2013). Also, as predicted by our our model, experimental manipulations have found that the relative fitness of cheats is higher when they are at lower frequencies in the populations, and at higher densities (frequency and density dependent selection) (Raymond et al., 2012).

Our model also makes novel testable predictions. We predicted that the fitness of cells that do not produce toxin (cheaters) depends upon an interaction between aggregation and density (fig. 4), and that toxin production should peak at intermediate group sizes (fig. 1). These predictions could be tested with field manipulations or experimental evolution. Our results also suggest the possibility for interactions between evolutionary and ecological (population) dynamics, that require further theoretical and empirical work. For example, low cell cell densities at the start of a season would favour cells that produce toxin (cooperators), which would lead to an increase in cell densities. This would favour cells that do not produce toxin (cheaters), which could reduce cell densities and now favour toxin producers again. Furthermore, these changes in cell densities and the frequency of toxin producers would also impact on the population dynamics of their invertebrate hosts, which could also influence the number of cells infecting each host (Raymond et al., 2012). These dynamics could potentially lead to seasonal patterns and/or intermittent epidemics of *B. thurigiensis*. The interplay of evolutionary and ecological dynamics between toxin non-producers and toxin producers has previously been demonstrated over the production of an enzyme to beak down sucrose in yeast (Gore et al., 2009; Sanchez and Gore, 2013).

**Figure 4:**
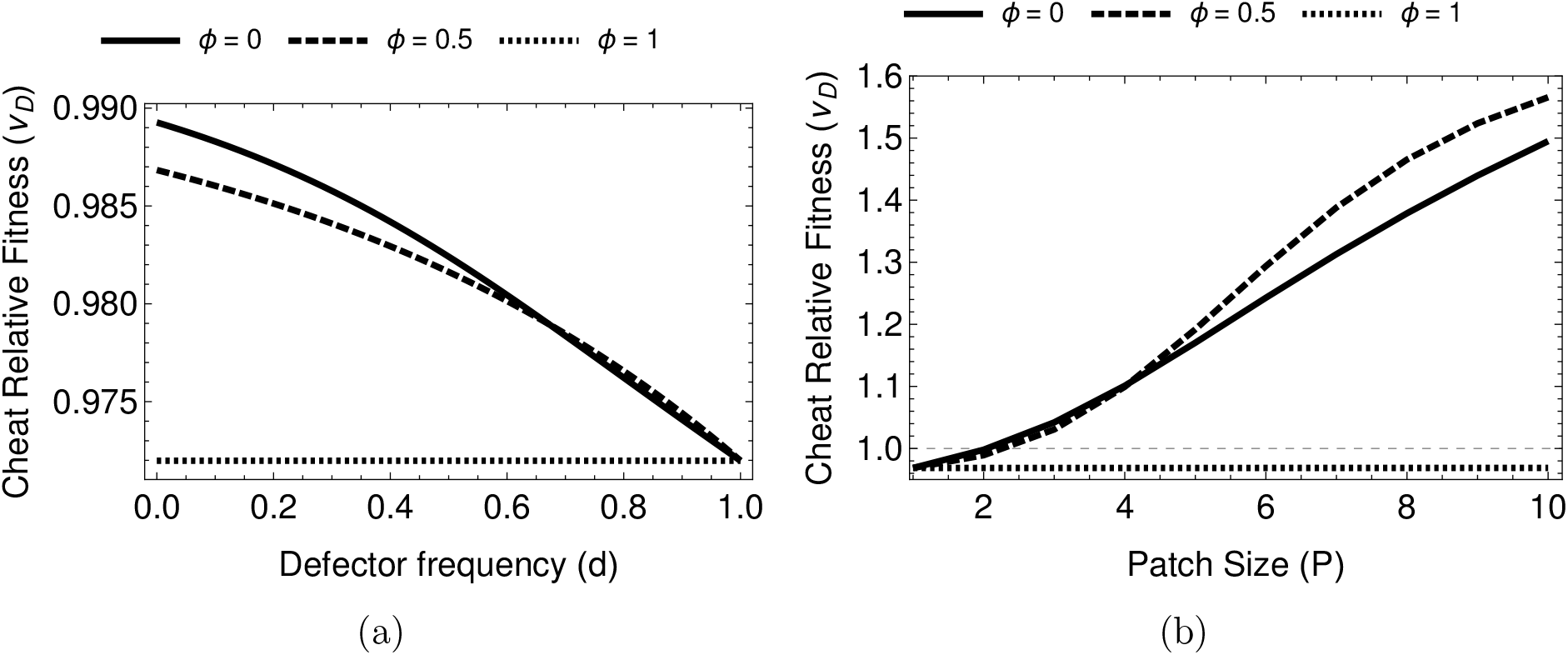
**(a)** Frequency dependence is still present in the base case of no aggregation as aggregation increases a critical point is reached at full aggregation where frequency dependence disappears **(b)** Increasing density increases cheat fitness as long as aggregation is again less than one. At full aggregation the density dependent effect disappears.

**Figure 5:**
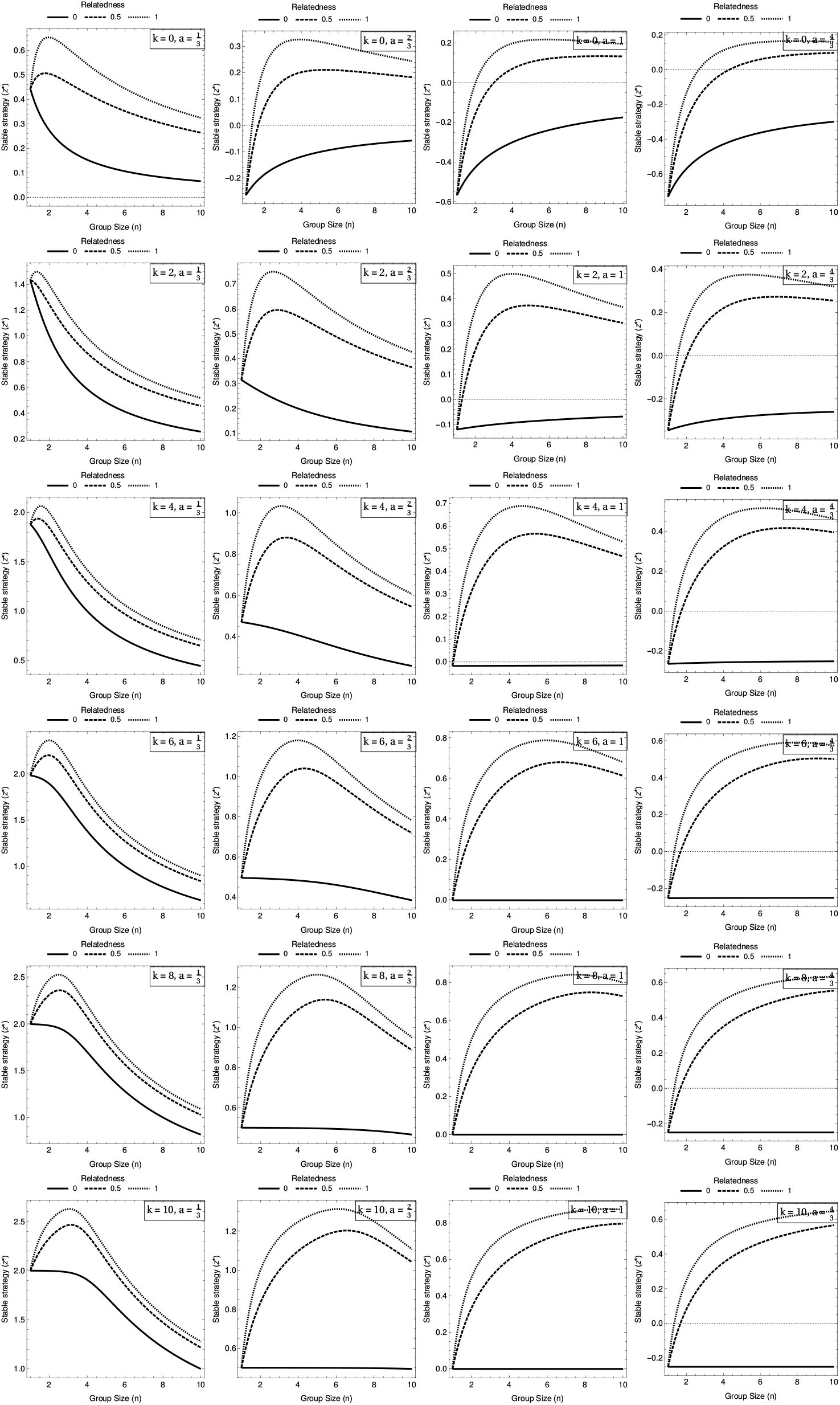

**Figure 6:**
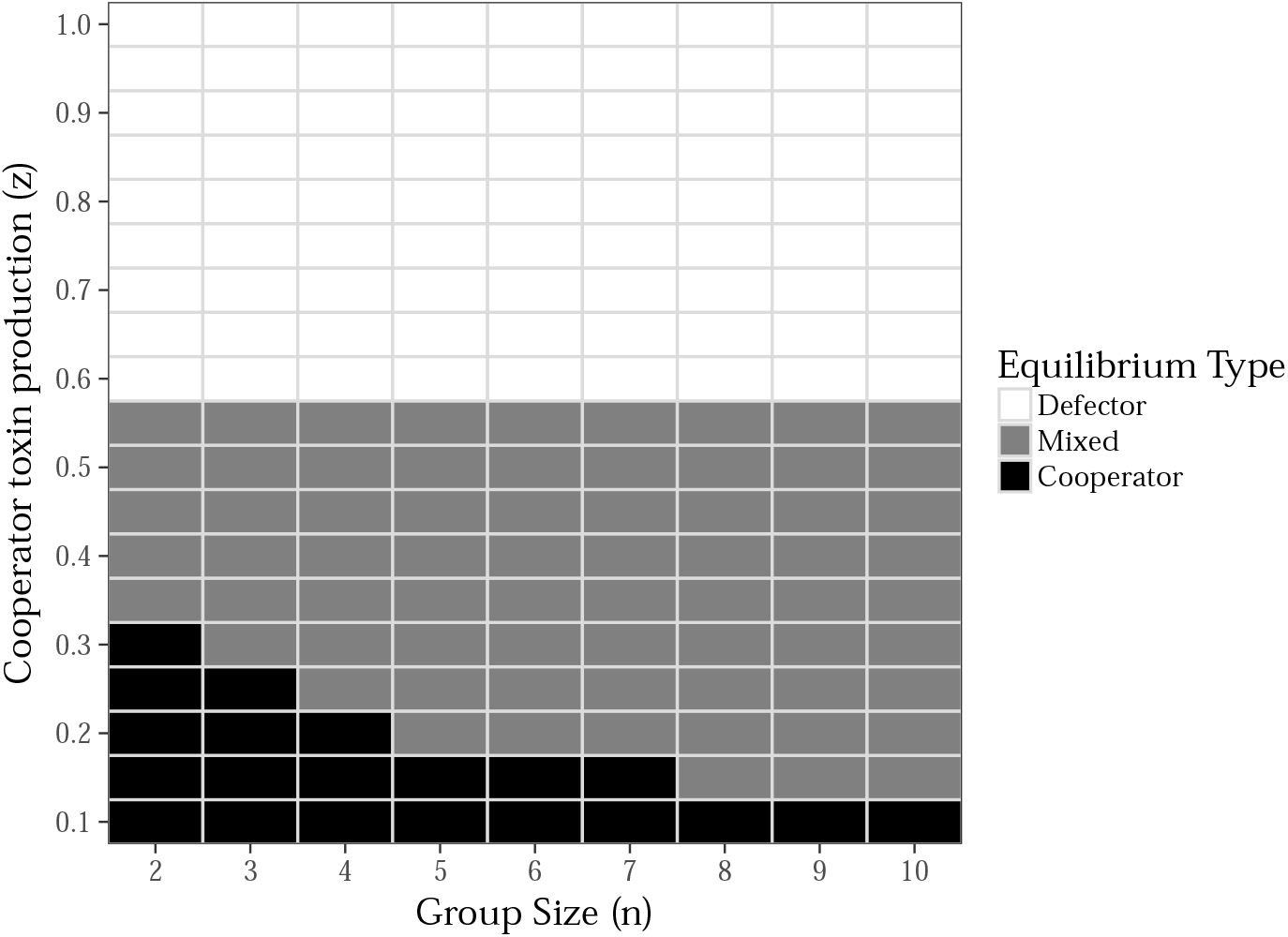
This figure shows the dynamics of a population of cooperators and defectors as described by eq. (10). Each point represents a poulation with *n* group size and cooperators that produce *z* toxin. Using the criteria for the gain sequence for each population we cklassify it as either a defector only equilibrium a cooperator only one or a mixed equilibrium where the two strategies coexist. The graph was drawn using the parameters, 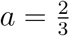
and *k* = 2

## A Uninvadibility condition

We construct a measure of the change in inclusive fitness caused by changing the focal actors strategy. This is done by expanding the total derivative of the fitness function with respect to a dummy variable for the underlying gene, which yields the expanded derivative (Taylor and Frank, 1996):

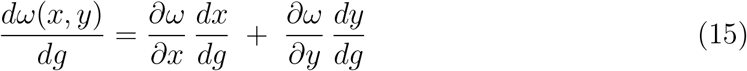

Following from Taylor and Frank (1996) we make the substitutions of the phenotypic derivatives for regression coefficients and then simplify to get an expression of change in inclusive fitness:

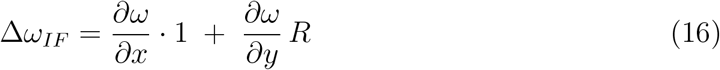

In the paper we then analyse the behaviour of ∆*ω_IF_* and 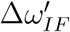 to characterise the equilibrium as maximal and convergent. Cooper and West (2018) method is used to determine if the equilibrium is unavailable. In brief, we consider the second derivative of the total derivative taken to obtain the inclusive fitness effects (Taylor and Frank, 1996). This expands into a long chain rule where we drop all higher order terms (*∂g*^2^ etc.) as negligible and substitute the regression coefficients as before, leaving us with:

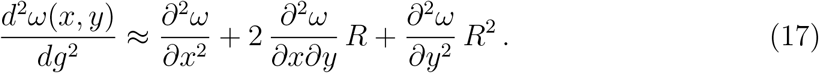

When this expression is less than zero we can say the equilibrium found is uninvaidable.

## B Sign of the ProductLog

*W* is a Lambert-W function which is strictly positive. The Lambert-W function or ProductLog function is the inverse of the functions in form *Xe^X^*:

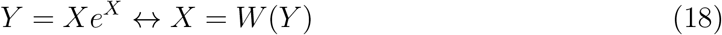

In this case the function in full is:

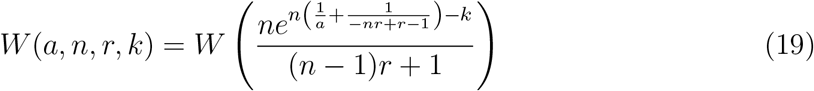

From the above we can see that — assuming: *a* > 0, 0 ≤ *r* ≤ 1, *n* ≥ 1, *k* ≥ 0 — then the function within the brackets will be positive and therefore the value of the function will be a positive real number.

## C Analysis of the effect of relatedness

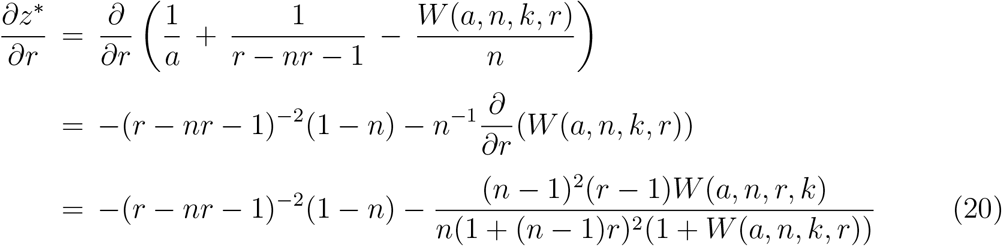

The expression obtained in eq. (20) is indeed always greater than or equal to zero forall values of r in the internal [0, 1]. We can see this by first remembering that the function *W* is always positive for any parameter set which is biologically reasonable — *a* > 0, 0 ≤ *r* ≤ 1, *n* ≥ 1, *k* ≥ 0. We then see that the first term is positive in the denominator (a squared term) and negative or zero in the numerator (1 − *n* where *n* ≥ 1), negating a negative is of course positive so the first term has positive effect on slope. The second term is also negative in the numerator as *r −* 1 is always zero or negative. Again it has a positive effect as it is negated from the slope. Therefore as long as the parameters are biologically reasonable the effect of increasing *r* is to increase investment in the toxin an individual produces.

## D Parameter sweep for the effect of group size

In the paper we assume a cost of two-thirds and a threshold of two in all scenarios. This was done so that in the case of two individuals total investment by both is necessary to reach the threshold value in *β*(*z*). The reason that the cost was set to 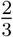 was to represent the fact that the tradeoff is against future investment not current investment. In fig. 5 we can see a greater range of parameters which are presented here to show that the patterns found are generally true across a reasonable range of parameter space.

## E Gain function and interior rest points

A property of the payoff *ω*(*z, y*) (eq. (10)) is that it is a polynomial in Bernstein form (Peña et al., 2014, 2015). This allows us to draw general conclusions about the shape and behaviour of this function by looking at a simple gain function. In essence we can calculate *a_i_* as the payoff for cooperating when *i* others cooperate, and *b_i_* as the payoff when defecting and *i* others cooperate.

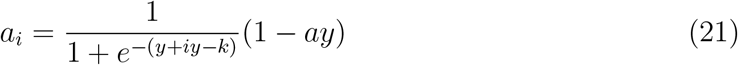

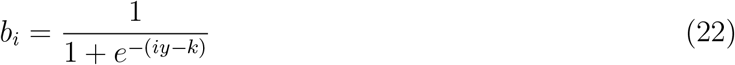

These are used to generate a measure of the gain from switching given *i* cooperators:

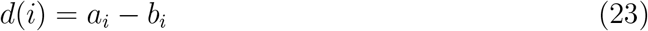

Which gives a gain sequence:

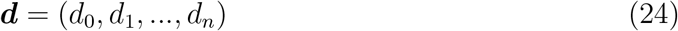

Now the purpose of this process is that the signs of the elements in the gain sequence, ***d***, tell us the stability of the two trivial rest states of the system and the number and stability of any interior rest points; assuming evolution occurs in an infinitely large well-mixed population (Peña et al., 2014). We are interested in three properties of the sequence:

1. If the sign of the first element (*d*_0_) is negative then the rest state of full defection is stable.
2. If the last element of the sequence (*d_n_*) is positive then the rest point of full cooperation is stable.
3. If there is one sign change in the sequence then there exists a unique interior rest point, furthermore, if the first element is positive then both trivial rest cases (all defect, all cooperate) are unstable and the interior point is stable.

Figure 6 shows the case with the parameters: 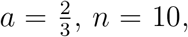 and *k* = 2. There exists a parameter range for the cooperating strategy between *z* = (0.1, 0.5] where there is a stable interior rest point — cooperators and defectors co-exist. When the trait is sufficently low then there is stable point when the population is all cooperators and when the trait value is higher than 0.5 the only stable scenario is all defectors. From the equilibrium game before we might expect the toxin production value to be around *z\** = 0.107 (3s.f.). This gives an initial element to the gain vector of, *d*_0_ = 0.00237 (3s.f.), and a final element of, *d*_0_ = 0.000457 (3s.f.), with no sign change in between. This indicates that the ESS solution to the static game would give a fully cooperative equilibrium in the dynamic game given these parameters.

